# gcaPDA: A Haplotype-resolved Diploid Assembler

**DOI:** 10.1101/2021.05.31.446328

**Authors:** Min Xie, Linfeng Yang, Chenglin Jiang, Shenshen Wu, Cheng Luo, Xin Yang, Lijuan He, Shixuan Chen, Tianquan Deng, Mingzhi Ye, Jianbing Yan, Ning Yang

## Abstract

Generating chromosome-scale haplotype resolved assembly is important for functional studies. However, current *de novo* assemblers are either haploid assemblers that discard allelic information, or diploid assemblers that can only tackle genomes of low complexity. Here, we report a diploid assembler, gcaPDA (gamete cells assisted Phased Diploid Assembler), which exploits haploid gamete cells to assist in resolving haplotypes. We generate chromosome-scale phased diploid assemblies for the highly heterozygous and repetitive genome of a maize F_1_ hybrid using gcaPDA and evaluate the assembly result thoroughly. With applicability of coping with complex genomes and fewer restrictions on application than other diploid assemblers, gcaPDA is likely to find broad applications in studies of eukaryotic genomes.

## Background

Deciphering the genetic blueprint of a species is of fundamental importance for related researches. Nowadays, genome sequence with quality on par with, if not superior to, the human reference genome could be easily generated with long read sequencing [1, 2] and assistant approaches [3, 4]. Most genomes of the plant and animal species are diploid, however, current long read *de novo* assemblers [5-7] are mainly aiming to generate high-contiguity haploid mixed-haplotypes assembly. In haploid mixed-haplotypes assembly, homozygous or low heterozygous regions are collapsed into a single mixed haplotype, whereas highly heterozygous regions are assembled into separate contigs and the allelic contigs (haplotigs) would be removed. Therefore, haploid mixed-haplotypes assembly can’t fully represent the complete genetic blueprint of diploid species[8].

To resolve haplotypes, *de novo* assemblers such as FALCON-unzip[9] and hifiasm[10] try to build diploid assembly by distinguishing long reads of different haplotypes based on heterozygous single nucleotide polymorphisms (SNPs). This strategy requires reads error rate to be much lower than genome heterozygous rate. In addition, heterozygous SNPs are not evenly distributed along chromosomes, with long stretch of homozygous region scattered in the genome. When long reads fail to span adjacent heterozygous SNPs, haplotype phasing across this region can’t be correctly inferred and lead haplotype switches. Therefore, supplementary data that can assist in long-range phasing is imperative to achieve chromosome-scale haplotype construction. In this regard, parental whole genome shotgun reads (WGS) data [11], Hi-C data [12], Strand-seq data [13] and gamete cell data [14, 15] have been used to bridge adjacent haplotype blocks.

Trio binning[11] uses WGS data of two parents to partition the progeny’s long reads and then assembles paternal and maternal genome respectively. It can provide entirely phased diploid assembly, except for novel mutations that unique to the progeny’s genome or heterozygous variants that exist in the trio samples. However, parental samples are not always available, especially in plant and animal studies. Diploid *de novo* assemblers such as DipAsm [12], PGAS (trio-free Phased Diploid genome Assembly using Strand-Seq) [13], Gamete-binning [14] share a common assembly framework (**Supplementary Figure 1**), which involving building an initial assembly, calling and phasing SNPs using sequencing reads, partitioning long reads based on phased SNPs and assembling each haplotype genome separately with partitioned long reads. This framework has gain success in genomes of low heterozygosity. However, it is not well suited for species with highly heterozygous genomes, which are quite prevalent scenarios in nature. Highly heterozygous genomes pose great challenge to haploid assembler and usually result in low quality initial assembly with fragmented contigs, abundant mis-assemblies and haplotigs. It is difficult to remove haplotigs thoroughly, while retaining haplotigs in the initial assembly will mess with SNP calling[16] and phasing process and generate poor haplotype-resolved assembly. Hence, it is particularly urgent to develop a diploid assembler that can cope with highly heterozygous genomes.

In this study, we report a diploid assembler, gcaPDA (gamete cells assisted Phased Diploid Assembler), that can generate chromosome-scale phased diploid assemblies for highly heterozygous and repetitive genomes using PacBio HiFi data, Hi-C data and gamete cell WGS data. gcaPDA offers equivalent performance to the trio-dependent method, with 98% phasing accuracy and >99% genome completeness. Both of the reconstructed haplotype assemblies generated using gcaPDA have excellent collinearity with their corresponding reference assemblies. Additionally, structural variations between reference genomes, including inversions and indels, are well-resolved. Having demonstrated its utility with a large and complex genome, it is plausible that gcaPDA can be easily applied to most diploid eukaryotic species and may find broad application in the coming diploid genome era.

## Results

### Schematic of the gcaDPA assembler

We have developed a diploid assembler, gcaPDA, to generate chromosome-scale phased genome assembly for diploid species. gcaPDA requires PacBio HiFi data and Hi-C data of an individual. In addition, short read WGS data of haploid gamete cells from the same individual are required to assist in long-range phasing. As illustrated in **Fig. 1** and **Supplementary Figure 2**, gcaPDA consists of 4 major steps: 1) generating an initial assembly; 2) reconstruction of haplotypes; 3) partition and normalization of gamete cell reads and 4) generating chromosome-scale phased diploid assembly.

**Figure 1.**
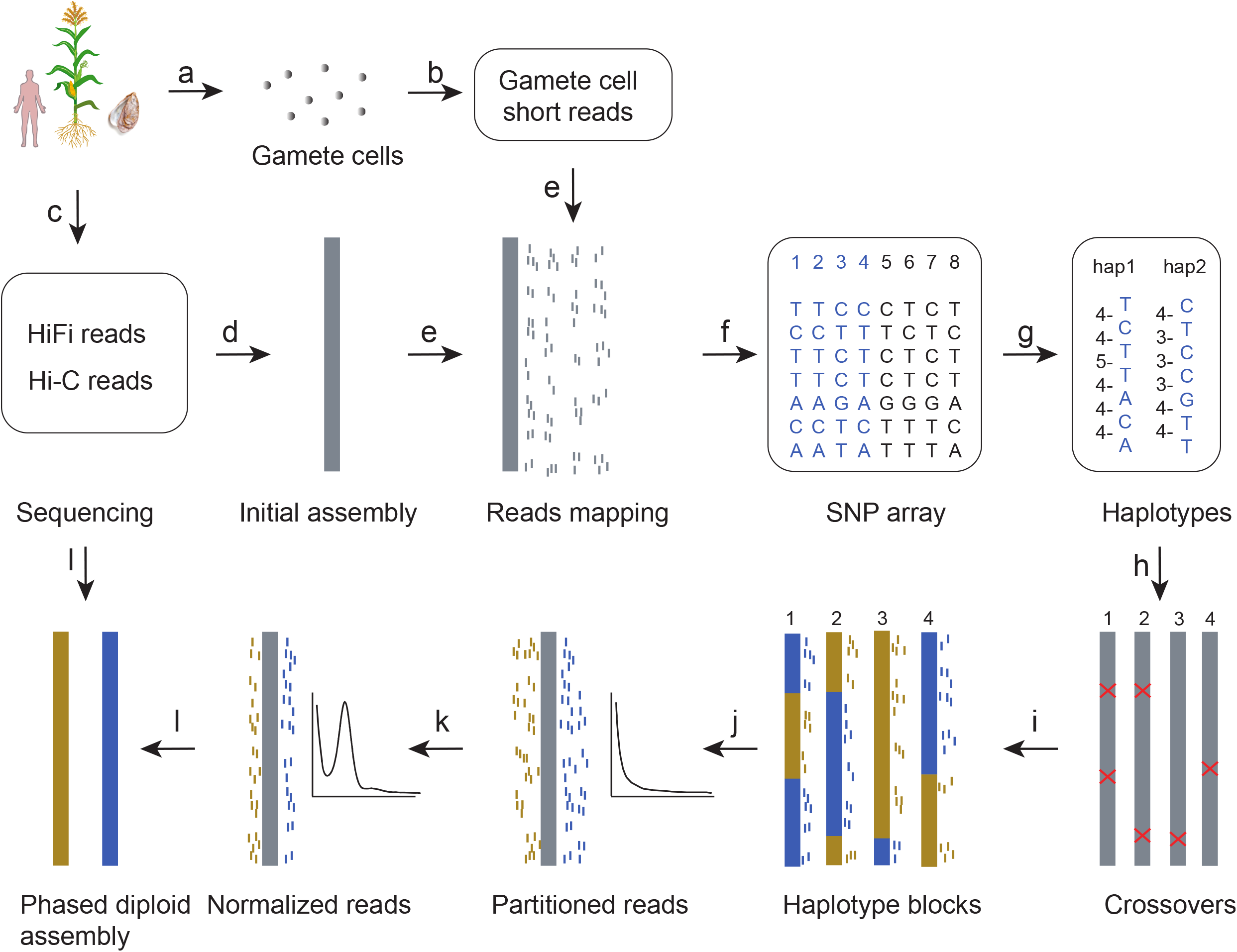
Schematic of gcaPDA workflow. **a)** Gamete cells (N=40) are isolated from focal individual **b)** Whole genome shotgun sequencing is performed to generate gamete cell short reads. **c)** HiFi reads and Hi-C reads are generated by sequencing somatic tissues. **d)** An initial assembly is generated by assembling HiFi reads into contigs and scaffolding contigs into superscaffolds with Hi-C data. **e)** Short reads of gamete cells are then mapped to the initial assembly and **f)** SNPs were identified for each gamete cells. **g)** Chromosomal-scale haplotypes of the sequenced individual are reconstructed based on gamete cell SNP arrays using major voting strategy^23^, with number of gamete cells that supports adjacent SNP combination were shown on the left. By comparing SNPs of each gamete cell with reconstructed haplotypes, **h)** crossovers and **i)** haplotype blocks of gamete cells can be determined. Gamete cell reads are **j)** partitioned based on haplotype blocks and **k)** normalized by k-mer depth to mimic genome coverage distribution of regular parental WGS reads. At last, **l)** HiFi reads and partitioned normalized gamete cell reads are then used to construct phased contigs using hifiasm^11^ and scaffolded into phased pseudochromosomes with Hi-C data.

### Generating sequencing data for a maize F_1_ hybrid

Maize has a very representative large and complex genome[17], which is suitable for testing the performance of gcaPDA. We selected a maize F_1_ hybrid derived from the crossing between two inbred lines B73 and SK as test sample.

High quality reference genomes are available for both parents (B73[18] and SK[19]) which could be used to benchmark the phased diploid assemblies. In total, 126 Gb and 129 Gb PacBio HiFi reads, with an average read length of 14Kb, were simulated based on published genome sequences of B73[18] and SK[19], respectively (**Supplementary Table 1**). Combining simulated B73 and SK PacBio HiFi reads together represents the sequenced reads of the F_1_ hybrid. K-mer analysis shows that the heterozygosity, haploid genome size and repeat content of the F_1_ hybrid genome are 1.98%, 2.16 Gb and 78% (**Supplementary Figure 3**), respectively, which indicates it’s a large, highly heterozygous and repetitive genome.

To assist chromosome-scale assembly, 40 microspores (hereafter referring to as gamete cells) were isolated from 10 tetrads of the F_1_ hybrid (**Fig. 1a**). DNA was extracted from each gamete cell and followed by multiple displacement amplification (MDA)[20] to generate enough DNA for library construction and sequencing[21] (**Fig. 1b**). Around 45 Gb (∼20-fold) high quality short reads were generated for each gamete cell (**Supplementary Table 3**). In addition, 427 Gb Hi-C data were also generated from root tissues of the F_1_ hybrid (**Fig. 1c** and **Supplementary Table 2**).

### Generating an initial assembly for the F_1_ hybrid

The simulated HiFi reads were assembled into contigs using haploid assembler FALCON[9] (**Fig. 1d**). The FALCON assembly is comprised of 2,903 Mb primary contigs and 685 Mb alternative contigs, with an contig N50 of 1.3 Mb (**Table 1**). The total length of primary contigs was larger than both the reference genomes of SK (2,161Mb) and B73(2,106 Mb), indicating there are ∼742 Mb haplotigs in the primary contigs. Of these, 432 Mb haplotigs can be tagged by purge_haplotigs[16], while the remaining 310 Mb haplotigs are undetectable (**Supplementary Table 4**). Since haplotigs in the primary contigs are hard to remove thoroughly, we choose to keep all of them and generated an initial assembly by scaffolding all the primary contigs into super-scaffolds with Hi-C data (**Fig. 1d**). The longest 10 supper-scaffolds (corresponding to 10 chromosomes) covers 97.5% of the initial assembly.

**Table 1.** Evaluation of FALCON assembly, Hifiasm assembly, Trio assembly and gcaPDA assembly.

| Assembly | Contigs |  |  | k-mer completeness (%) |  |  |  | Gene completeness (%) |  |  |  |
| --- | --- | --- | --- | --- | --- | --- | --- | --- | --- | --- | --- |
|  | Total (Mb) | No. | N50 (Mb) | All | Hap SK | Hap B73 | PPV | Comp | Dup. | Frag. | Mis. |
| **Reference SK** | 2,154 | 671 | 223.2 | 76.15 | 100.00 | 0.00 | NA | 96.1 | 6.0 | 1.3 | 2.6 |
| **B73** | 2,104 | 267 | 220.8 | 75.97 | 0.00 | 100.00 | NA | 95.5 | 6.1 | 1.5 | 3.0 |
| **B73 + SK** | 4,258 | 938 | 223.2 | 100.00 | 100.00 | 100.00 | NA | 96.9 | 94.8 | 0.9 | 2.2 |
| **FALCON** |  |  |  |  |  |  |  |  |  |  |  |
| Pri | 2,903 | 5,014 | 1.7 | 85.69 | 73.92 | 72.56 | NA | 78.2 | 11.0 | 6.1 | 15.7 |
| Alt | 685 | 9,221 | 0.2 | 32.10 | 19.68 | 20.96 | NA | 35.3 | 1.0 | 4.5 | 60.2 |
| Pri + alt | 3,589 | 14,235 | 1.3 | 95.25 | 91.91 | 91.69 | NA | 83.1 | 39.2 | 5.2 | 11.7 |
| **Hifiasm** |  |  |  |  |  |  |  |  |  |  |  |
| Pri | 3,393 | 1,164 | 19.9 | 91.66 | 83.11 | 82.26 | NA | 96.6 | 48.5 | 0.6 | 2.8 |
| Alt | 871 | 3,708 | 2.8 | 36.62 | 19.00 | 18.11 | NA | 49.3 | 3.7 | 1.8 | 48.9 |
| Pri + alt | 4,264 | 4,872 | 14.6 | 99.96 | 99.91 | 99.91 | NA | 97.4 | 87.7 | 0.6 | 2.1 |
| **Trio** |  |  |  |  |  |  |  |  |  |  |  |
| HapSK | 2,156 | 688 | 82.0 | 76.24 | 99.70 | 0.45 | 99.55 | 96.5 | 5.8 | 0.8 | 2.7 |
| HapB73 | 2,113 | 334 | 75.6 | 76.11 | 0.73 | 99.78 | 99.27 | 95.9 | 6.5 | 1.2 | 2.9 |
| HapSK + hapB73 | 4,269 | 1,022 | 75.6 | 99.92 | 99.79 | 99.88 | 99.41 | 97.6 | 94.9 | 0.4 | 2.1 |
| **gcaPDA** |  |  |  |  |  |  |  |  |  |  |  |
| HapSK | 2,162 | 767 | 55.3 | 76.19 | 98.87 | 1.78 | 98.25 | 95.9 | 6.5 | 1.4 | 2.7 |
| HapB73 | 2,159 | 699 | 57.0 | 76.41 | 2.25 | 99.46 | 97.77 | 95.4 | 6.2 | 1.6 | 3.0 |
| HapSK +hapB73 | 4,320 | 1,466 | 55.3 | 99.67 | 99.06 | 99.56 | 98.01 | 96.9 | 94.5 | 1.0 | 2.2 |

\* PPV indicates positive predictive value

### Reconstruction of chromosome-scale haplotypes for the F_1_ hybrid

Short reads of gamete cells were mapped to the initial assembly and SNPs were identified for each gamete cell (**Figs. 1e** and **1f**). Nine out of the 40 gamete cells have abnormal SNP heterozygous rate or missing rate (**Supplementary Table 5** and **Supplementary Figure 4**), which could be caused by contamination of adjacent cells or insufficient genome coverage. These gamete cells were considered to be low-quality and excluded from downstream analyses. In general, 1,387,594 to 2,423,031 high confident SNPs were identified in each gamete cell (**Supplementary Table 5**).

Chromosome-scale haplotypes of the sequenced F_1_ hybrid were reconstructed based on the SNPs that located on 10 chromosomes using a major voting strategy with R package Hapi[22] (**Fig. 1g**). Totally, we obtained 2,721,839 phased SNPs in the reconstructed chromosome-scale haplotypes, which are evenly distributed across chromosomes (**Supplementary Figures 5 and 6**). Haplotype blocks in each gamete cell could be identified by comparing the genotype at each SNP locus with that of the reconstructed chromosome-scale haplotypes (**Figs. 1h, 1i and Supplementary Figure 6**).

### Partition and normalization of gamete cell short reads

Reads of each gamete cell that belonged to haplotype blocks of B73 or SK were extracted and merged separately (**Fig. 1j**). Reads of each haplotype were normalized by k-mer depth to mitigate the extremely uneven genome coverage caused by MDA procedure (**Fig. 1k**). After normalization, k-mer depth distribution of the haplotype reads is similar to that of parental WGS data (**Supplementary Figure 7)** and ready for use by trio-dependent *de novo* assemblers (HiCanu, hifiasm, etc.).

### Generating phased diploid genome assemblies for the F_1_ hybrid

The phased diploid genome assemblies were generated using hifiasm[10], with simulated HiFi reads and k-mers derived from normalized haplotype reads (**Fig. 1l**). In total, 2,162 Mb contigs were assigned to SK haplotype (HapSK contigs) and 2,159 Mb contigs were assigned to B73 haplotype (HapB73 contigs), with a contig N50 of 55.3 Mb and 57.0 Mb, respectively. The hapSK contigs and hapB73 contigs were further scaffolded into chromosomes with Hi-C data, respectively. This chromosome-scale phased diploid assembly, including both hapSK assembly and hapB73 assembly, is referred to gcaPDA assembly hereafter.

### Comparing gcaDPA with other methods

To compare the performance of gcaPDA with other methods, we generated a “Hifiasm assembly” with using hifiasm with only HiFi reads, and a “Trio assembly” using hifiasm with HiFi reads and simulated parental WGS reads. Genome assemblies were evaluated in three aspects: contiguity, completeness and accuracy.

Genome contiguity is usually measured with contig N50. Comparing to FALCON (contig N50: 1.3 Mb), diploid assembler hifiasm achieved better contiguity (contig N50: 14.6Mb) by resolving haplotypes with HiFi reads. After adding in supplementary data (gamete cell WGS data or parental WGS data), contig N50 were further improved in gcaPDA assembly (55.3 Mb) and Trio assembly (75.6 Mb) (**Table 1**). In summary, contiguity of heterozygous genome can be dramatically improved by resolving haplotypes.

Genome completeness were evaluated by k-mer, and BUSCOs[23] (Benchmarking Universal Single-Copy Orthologs), and assembled genome size. In total, 799 million distinct k-mers were counted in SK and B73 genomes. Of these, 52.1% were shared between B73 and SK, while 23.8% and 24.0% k-mers were unique to B73 genome (refer to as B73 hapmer hereafter) and SK genome (refer to as SK hapmer hereafter) (**Supplementary Table 6**). All assemblies achieved >99% k-mer completeness except for FALCON assembly which has a k-mer completeness of 95% (**Table 1**). Similarly, all assemblies achieved >99% hapmer completeness except for FALCON assembly which has a hapmer completeness of ∼92%. In addition, all assemblies achieved overall BUSCO completeness comparable to that of the parental genomes, except for FALCON assembly. In accordance with the evaluation by k-mer and BUSCOs, assembled genome size of all assemblies are comparable to that of the parental genomes, except for FALCON assembly (**Table 1**).

Phasing accuracy of genome assemblies were evaluated with hapmer. The overall phasing accuracy (positive predictive value, PPV) of the gcaPDA assembly is 98.01%, comparable to that of the Trio assembly (**Table 1**) and assemblies reported in recent studies [14, 24, 25]. PPV value was not calculated for FALCON assembly or Hifiasm assembly, since contigs in these two assemblies were not assigned to specific haplotypes. Instead, we investigated chimeric contigs that contain both SK and B73 hapmer. In FALCON assembly and Hifiasm assembly, there are many contigs contain hapmers from both haplotypes (**Fig. 2**). In comparison, only few contigs in gcaPAD assembly and Trio assembly contain hapmer from the other haplotype (**Fig. 2**). With hapmer, we also detected haplotype blocks from the other haplotype in the hapSK and hapB73 assembly of gcaPDA (**Fig. 3 and Supplementary Table 8**). It is worth to note that un-anchored sequences are more prone to be mis-assigned to the other haplotype, when comparing with sequences that anchored to chromosomes **(Supplementary Table 8**).

**Figure 2.**
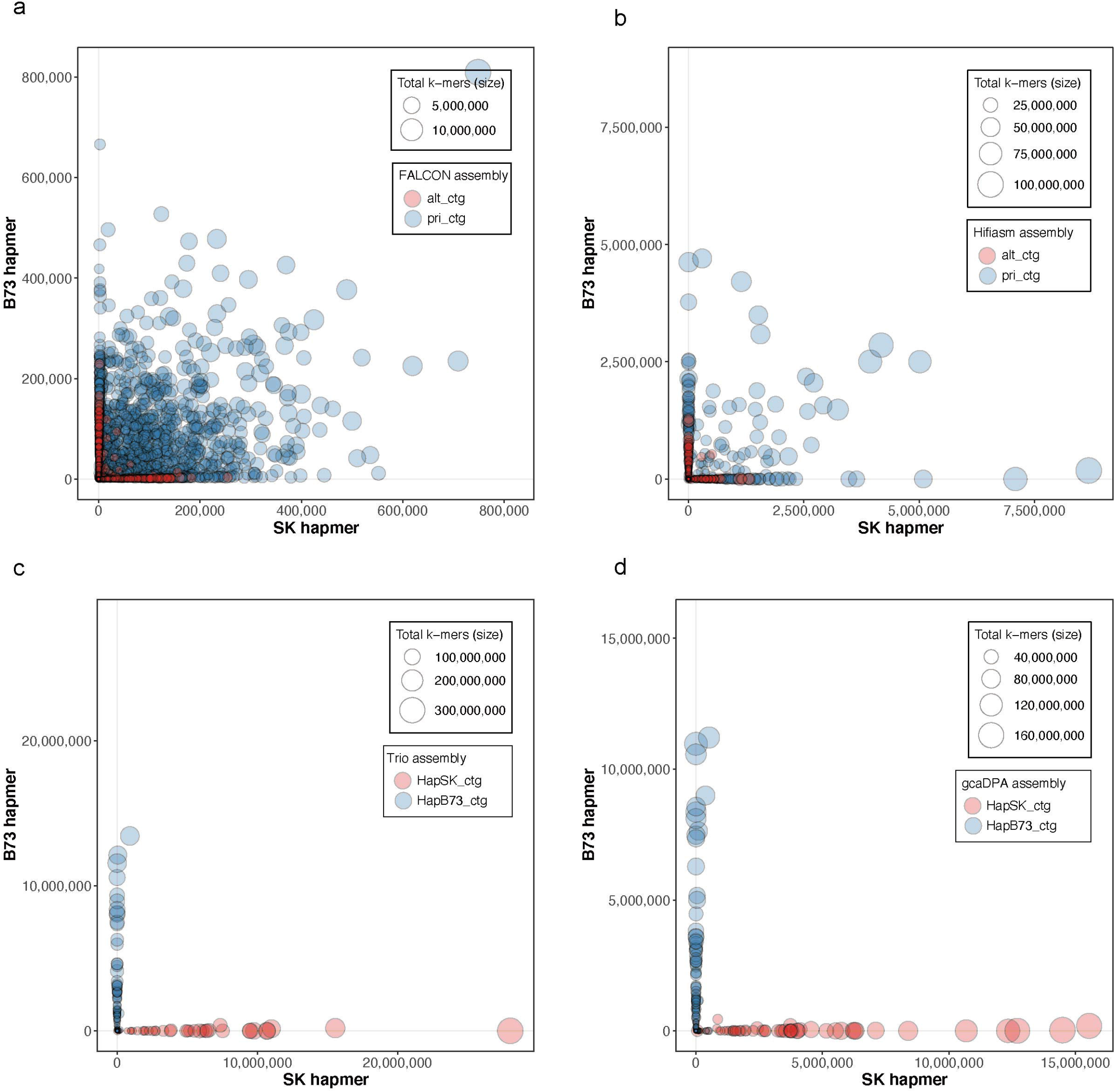
Comparison of phasing accuracy of contigs of different assemblies. Each contig is represented by a circle, with circles of primary/hapB73 contigs filled red and alternative/hapSK contigs filled blue. The size of a circle is proportional to the total number of k-mer in the contig. The x and y axes refer to the number of SK hapmer and B73 hapmer identified in a contig, respectively. Panel **a)** FALCON assembly, **b)** Hifiasm assembly, **c)** Trio assembly, **d)** gcaPDA assembly.

**Figure 3.**
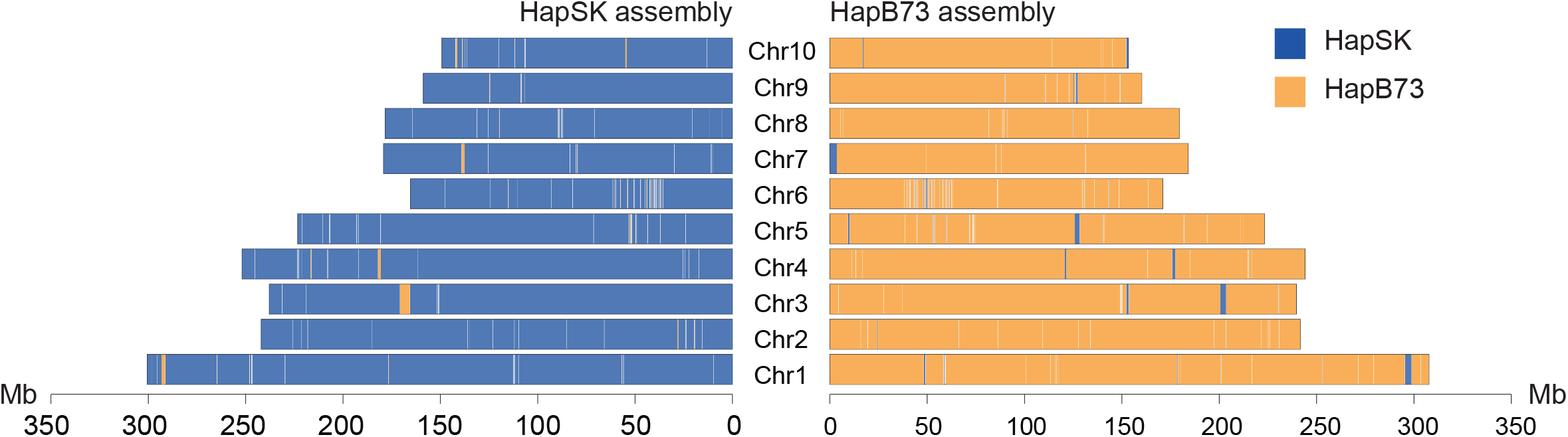
Haplotype blocks in the gcaPDA assembly. Each chromosome is represented by a rectangle, with width proportional to chromosome length. Haplotype blocks are defined by consecutive hapmer of the same haplotype.

### Comparing gcaDPA assembly with parental reference genomes

With reference genome sequences for both parents of the F_1_ hybrid available, accuracy of gcaPDA assembly was further evaluated by comparing with parental genomes. The HapSK and HapB73 assembly were compared with SK and B73 reference genomes, respectively. The hapSK assembly covers 99.04% of the SK reference genome, with an average alignment identity of 99.99%, while the hapB73 assembly covers 99.72% of the B73 reference genome, with an average alignment identity of 99.99% (**Supplementary Table 7**). In general, near perfect collinearity was observed between haplotype-resloved assembles and corresponding parental genomes (**Supplementary Figure 8**).

Notablely, gcaPDA could phase the structural variations between B73 and SK genome properly. For example, a large inversion (*c*. 8 Mb) and a large indel (*c*. 3 Mb) between B73 and SK genomes were correctly recovered in the hapSK and hapB73 assembly (**Fig. 4a**). In addition, we inspected the *ZmBAM1d* locus^19^ which is highly divergent between B73 and SK genomes. We found that the *ZmBAM1d* locus were also perfectly phased in the gcaPDA assembly, resulting in haplotype sequences identical to respective reference genome (**Fig. 4b**).

**Figure 4.**
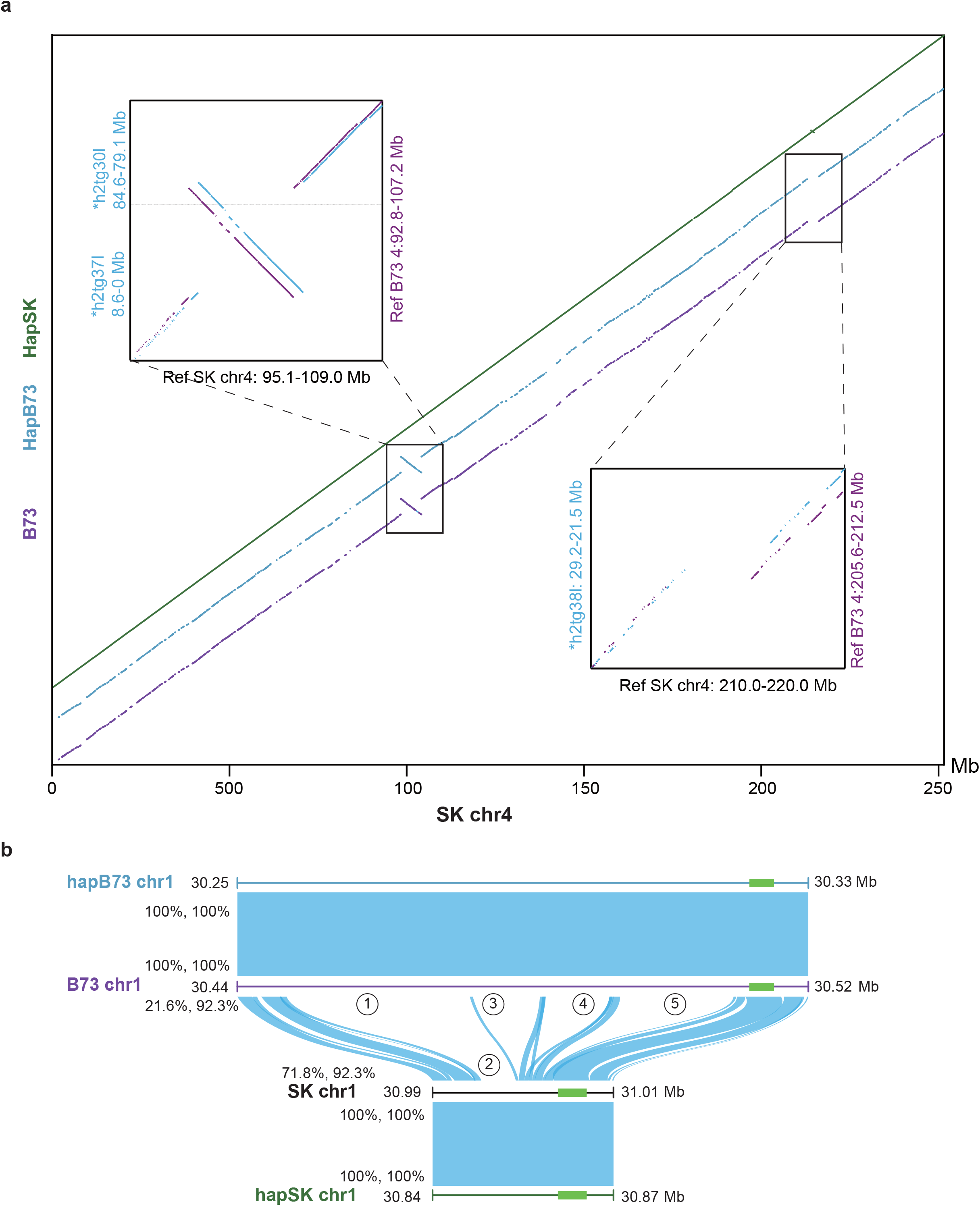
Sequence comparison of different assemblies. **a)** Chromosome 4 of HapSK, hapB73 assembly (generated by gcaPDA) and reference B73 (B73) assembly were aligned to chromosome 4 of reference SK assembly (the alignments colored with green, blue and purple, respectively). Inset at upper left: a large inversion (*c*. 8 Mb) between B73 and SK. Inset at lower right: a large indel (*c*. 3 Mb) between B73 and SK. **b)** Sequence alignments of the *ZmBAM1d* locus, which is known as highly divergent between B73 and SK. Numbers with circles indicate large indels between SK and B73. Coverage and identity of the alignments are shown on the left. *ZmBAM1d* genes is represented as a green brick.

## Discussion

Diploid genomes comprise two sets of homologous chromosomes that are slightly different from each other. Current diploid assemblers (such as DipAsm [12], gamete binning[14], PGAS[13]) are tailored for genomes of low complexity and not suitable for heterozygous genomes. In the present study, we developed gcaPDA, a gamete cell-based method which generates chromosome-scale haplotype-resolved diploid assembly for species with highly heterozygous genomes, thus providing access to the genetic variations present in both sets of homologous chromosomes of diploid cells.

In standard trio binning[11], parental data are used to resolve haplotypes. For a highly heterozygous individual, it is natural to assume that its parents are highly heterozygous, too. Notably, phasing information can be fuzzy at sites where all trio-samples have a heterozygous genotype[10]. And parent information is not always available, which limits the application of trio binning method. In contrast, gcaDPA does not require parental information. The gamete cells used to generate input data for gcaPDA are naturally haploid, and preserves unambiguous chromosome-scale phasing information. Thus, gcaPDA’s performance is not affected by the heterozygosity level of the sequenced individual.

In DipAsm [12], gamete binning[14], PGAS[13], and gcaDPA, an initial assembly is generated for SNP calling and phasing (**Supplementary Figure 1**). For highly heterozygous genome, the initial assembly may contain haplotigs, which lead to missing SNP calls (false negative)[16]. In addition, SNP calling at repetitive genomic regions are prone to generate false positive calls[26]. Both false positive and negative SNP calls can complicate SNP-based long reads partition in DipAsm, gamete binning, and PGAS methods. Furthermore, partition long reads before assembly process is not the recommended way of diploid assembly[10]. In contrast, gcaPDA partitions gamete cell reads based on haplotype blocks instead of individual SNPs, to increase the tolerance to potential false negative and positive SNP calls. In addition, gcaPDA used all the HiFi reads to construct assembly graphs, with haplotype-specific k-mer derived from gamete cell reads to assist in resolving graph, and generate both haplotype assembly simultaneously (**Supplementary Figures 1 and 2**). Furthermore, the input data required by gcaPDA were generated by standard wet-lab and sequencing approaches which are accessible to researchers. In contrast, gamete binning needs to sequence hundreds of thousands of gamete cell genomes by 10x sc-CNV sequencing^14^, while this service is no longer supported by 10x Genomics. The Hi-C data used by DipAsm were generated with four restriction enzymes based on a modified protocol with Arima-HiC kit^12^ to achieve uniform per-base coverage of the genome and maintain the highest long-range contiguity signal, while it may be difficult to generate Hi-C data of comparable quality for plant species. Generation of Strand-seq[27] data required by PGAS pose great challenge for non-human species, which limits the application of PGAS.

There are several factors that affect the performance of gcaPDA. First of all, HiFi read length. We simulated HiFi reads with 14Kb length, however, current Pacbio Sequel II can generate HiFi reads with average length longer than 20 Kb by size selection of DNA fragments during library construction. Longer read length have better chance to span adjacent heterozygous loci, hence could further improve phasing accuracy. Second, contiguity and accuracy the initial assembly. The genome of F_1_ hybrid is highly repetitive and heterozygous, which poses a great challenge for FALCON and results in abundant short contigs. Short contigs that couldn’t be scaffolded into chromosomes won’t be phased by gcaPDA, while short contigs wrongly placed during Hi-C scaffolding analysis resulting in mis-assemblies (false inversion, translocation) in the initial assembly[28], which in turn lead to chunks of gamete cells reads wrongly assigned to the other haplotype. Improving contiguity of the initial assembly with longer HiFi reads, or incorporating optical mapping data to correct mis-placed sequences[29] might mitigate these issues and improve the performance of gcaPDA. Third, number of gamete cells. In this study, 31 gamete cells (∼20X WGS data for each cell) were used by gcaPDA. According to k-mer coverage accumulation curve (**Supplementary Figure 9**), 20 gamete cells shall suffice. However, with more gamete cells sequenced, phasing errors introduced by mis-placed contigs shall be alleviated and higher phasing accuracy can be achieved by gcaPDA.

All in all, taking the assembly of a real large, highly heterozygous and repetitive maize F_1_ hybrid genome as the positive control, we proved that gcaPDA could generate high quality haplotype-resolved and chromosome-scale diploid assembly for diploid species. In contrast to other diploid assemblers, gcaPDA does not rely on paternal information, tremendous amount of gamete cells or special sequencing approaches. As a result, gcaPDA is likely to find broad applications in studies of eukaryotic genomes.

## Materials and Methods

### Simulating reads based on reference genomes

Reference genome sequences of *Zea mays* var. SK was downloaded from ZEAMAP database(http://www.zeamap.com/ftp/01_Genomics/Genomes/)[30], while reference genome sequences of *Zea mays* var. B73 was downloaded from Ensembl Plant database[31] (release 46). In order to lift the contiguity limit capped by the reference sequences, unambiguous bases (‘N’s) within sequences were removed to generate gapless sequences. Reads were simulated based on SK or B73 gapless sequences. For short reads simulation, wgsim (parameters: -e 0.01) from samtools package[32] (version 1.9) was used. pbsim[33] (version 1.0.4) was used to simulate PacBio HiFi reads with read length and quality score randomly sampled from the previously published PacBio HiFi data[34].

### Genome survey analysis

Genome survey analysis was performed to profile features of the F_1_ hybrid genome. Simulated SK and B73 HiFi reads were broken into k-mer and then counted by Jellyfish[35] (version 2.2.10, k=21). Genome features such as haploid genome size, heterozygosity, repreat content were estimated using genomescope[36] (version 1) with k-mer frequencies outputted by Jellyfish.

### Hi-C library for maize F_1_ hybrid

Young root tissues of F_1_ hybrid (B73 x SK) were harvested. Hi-C proximity libraries were constructed by the previously described method with restriction enzyme MboI[4]. The libraries were size-selected to retain 350 bp DNA fragments and sequenced on MGISEQ2000 platform (MGI-Tech) to generate 150 bp paired-end reads.

### Single cell isolation, DNA extraction, amplification and sequencing

*Zea mays* var. SK was crossed with *Zea mays* var. B73 were crossed to generate seeds of F_1_ hybrid individuals. The seeds of F_1_ hybrid were planted and then immature tassels were harvested before they had emerged. Gamete cells (microspores) were isolated from tetrads as described in previous study[21]. DNA were extracted from each gamete cell using QIAGEN REPLI-g Single Cell Kit (Cat No. 150343), followed by multiple displacement amplification (MDA)[20] procedure to generate enough DNA for downstream experiments. A sequencing library was constructed for each gamete cell and then to be sequenced on MGISEQ2000 platform (MGI-Tech) to generate 150 bp paired-end reads.

### Generating initial assembly

Simulated SK and B73 HiFi reads were assembled into contigs using falcon[9] (version 1.4.4) from pb-assembly packages (https://github.com/PacificBiosciences/pb-assembly#citations), with parameter settings:

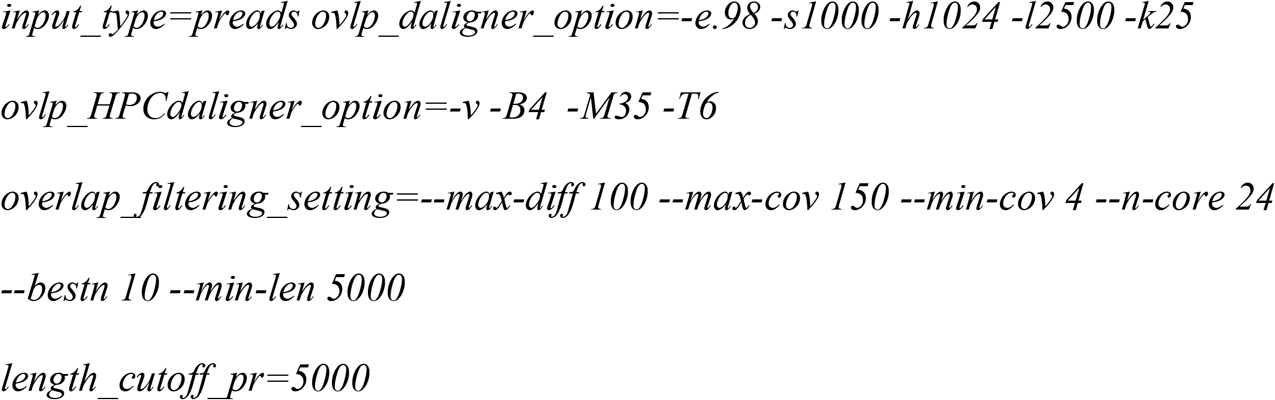

This assembly was referred to as FALCON assembly in the main text.

To link contigs to superscaffolds, Hi-C scaffolding analysis was performed. Briefly, Hi-C reads were pre-processed using HiC-Pro program[37] (v2.11.1) and then mapped to contigs using BWA[38] (v0.7). Low quality mapping (MAPQ=0) and duplicates were removed and Hi-C contract matrices were calculated using Juicer[39] (v1.6.2). The Hi-C contact matrices were fed to the 3D-DNA pipeline[28] (v180922, parameters: -m haploid -r 0) to order and orient contigs. Potential mis-joins were corrected manually to generate superscaffolds. The longest 10 superscaffolds were considered as pseudochromosomes of the initial assembly.

### Reconstruction of haplotypes and gamete cell reads partition

The raw reads of gamete cell were preprocessed to filter adapter sequences and low-quality reads using bbduk.sh (parameters: ktrim=r k=17 mink=7 hdist=1 tpe tbo qtrim=rl trimq=15 minlength=80) from BBTools package (https://sourceforge.net/projects/bbmap/). Reads that passed quality filtering were then mapped to the initial assembly using bowtie2[40] (version 2.3.5.1), with parameters “-X 800”. Single nucleotide polymorphisms (SNPs) were identified using bcftools[41] ‘mpileup’ (version 1.8,parameters: -d 500 -q 10 -- ff SECONDARY), followed by bcftools ‘call’ (parameters: -Ob -cv -p 0.01) implementation. Only bi-allelic SNPs, with quality value >=20 and allele frequency between 0.3 and 0.7 were selected. SNPs that located adjacent to (< 5 bp) indels were also filtered. Furthermore, SNPs in each cell sample with supporting reads depth <5 were replaced with ‘NA’ and treated as missing calls. Heterozygous rate and missing rate were calculated for each cell based on SNP calls. Since we sequenced haploid single cells, most of the SNPs identified in each cell should be homozygous, with lots of missing calls due to insufficient read coverage. Cells with abnormal level of heterozygous rate of SNPs (>5%) or missing call rate (<30% or >70%) indicate contamination or insufficient sequencing coverage, hence were considered as low-quality cells and excluded from downstream analyses.

The paternal and maternal haplotypes were reconstructed based on SNP array of qualified cells using Hapi[22]. By comparing genotypes of each gamete cell with reconstructed parental haplotypes, haplotype blocks can be identified for each cell. Gamete cell reads that mapped to haplotype blocks with the same parental origin were extracted and merged. Due to MDA procedure, the read depth of each cell is unevenly distributed across the genome. To mitigate this issue, merged reads of each haplotype were normalized by k-mer depth using BBnorm.sh (https://sourceforge.net/projects/bbmap/), to mimic regular whole genome sequencing (WGS) reads. The normalized short reads were then used as parental WGS data by diploid *de novo* assembler.

### *De novo* genome assembly

The normalized short reads of gamete cells of each haplotype were broken into k-mers using yak (https://github.com/lh3/yak), respectively. Simulated HiFi reads (B73 and SK), together with parental k-mers, were *de novo* assembled using hifiasm[10] to generate phased diploid genome assembly. The haplotype-resolved contigs were linked into superscaffolds using Hi-C reads, with methods described above. The final results of gcaPDA were two haplotype-resolved chromosome-level assemblies: hapSK assembly and hapB73 assembly. This final assembly was referred to as gcaPDA assembly in the main text.

In the same time, another haplotype-resolved diploid assembly were generated with HiFi reads (B73 and SK) and simulated parental WGS short reads using hifiasm. This assembly was referred to as Trio assembly.

Furthermore, a pseudo-haplotype assembly was generated using hifiasm, with only simulated HiFi reads. This assembly was referred to as Hifiasm assembly.

### Evaluation of genome assemblies

We broke the gapless reference genomes of B73 and SK into k-mer using meryl utility from Merqury package[42] (release 20200430). Total k-mer set and haplotype-specific k-mer set (hapmers) were computed based on B73 k-mer set and SK k-mer set. Genome completeness of each assemblies was evaluated with total k-mer set and phasing accuracy were evaluated with hapmers using Merqury package[42].

Gene completeness of the assemblies was evaluated using BUSCO[23] (version 3.0.2, lineage setting: embryophyta_odb9).

Whole genome sequence comparison between assemblies and reference genomes were performed using nucmer and visualized using mummerplot from MUMMER package[43] (version 4.0). Coverage and identity of the alignments were calculated using dnadiff implementation from MUMMER package. Only alignments span >10Kb were counted.

### Tagging haplotigs from FALCON primary contigs

The simulated SK and B73 HiFi reads were mapped to FALCON primary contigs using minimap2[44] with parameters “-ax asm20 --secondary=no”. Haplotigs in the FALCON primary contigs were tagged by purge_haplotigs[16] based on HiFi read coverage of each contig and the syntenic relationships between contigs.

## Declarations

### Authors’ contributions

J-B.Y., N.Y. and L-F.Y. designed the project. C-L.J., N.Y., C.L., S-S.W. contributed to sample preparation and wet-lab experiments. M.X., L-F.Y., X.Y., L-J.H., S-X.C. T-Q.D. and M-Z.Y. performed all data analysis. M.X. wrote the first draft of this manuscript. M.X., N.Y., and J-B.Y. wrote the final draft.

### Availability of data and materials

The data reported in this study are available in the CNGB Nucleotide Sequence Archive (CNSA: https://db.cngb.org/cnsa; accession number CNP0001702). The code of gcaPDA are available at https://github.com/BGI-shenzhen/gcaPDA.

## Acknowledgements

Not applicable.

## Ethics approval and consent to participate

Not applicable.

## Consent for publication

Not applicable.

