## Supplemental files for "gcaPDA: A Haplotype-resolved Diploid Assembler"

**Supplementary Figures**

Supplementary Figure 1. Comparison of framework of different diploid assemblers.

Supplementary Figure 2. The analysis workflow of gcaPDA.

Supplementary Figure 3. Genome survey of the SK x B73 F_1_ hybrid.

Supplementary Figure 4. Quality control of gamete cells based on SNP statistics.

Supplementary Figure 5. Distribution of distance between adjacent SNPs in the reconstructed haplotypes.

Supplementary Figure 6. Visualization of haplotype blocks of gamete cell S2.

Supplementary Figure 7. K-mer distribution of haplotype reads before and after normalization.

Supplementary Figure 8. Collinearity relationship between gcaDPA assembly and parental genomes.

Supplementary Figure 9. Accumulated percentage of genomic k-mers covered in gamete cells reads.

**Supplementary Tables**

Supplementary Table 1. Statistics of simulated reads.

Supplementary Table 2. Statistics of Hi-C sequencing reads.

Supplementary Table 3. Statistics of sequencing reads of gamete cells.

Supplementary Table 4. Purge haplotigs from FALCON primary contigs.

Supplementary Table 5. Statistics of SNPs identified for gamete cells.

Supplementary Table 6. Statistics of k-mer in the parental genomes.

Supplementary Table 7. Statistics of whole genome sequence comparison between haplotype assemblies and reference assemblies.

Supplementary Table 8. Statistics of haplotype blocks from the other haplotype.

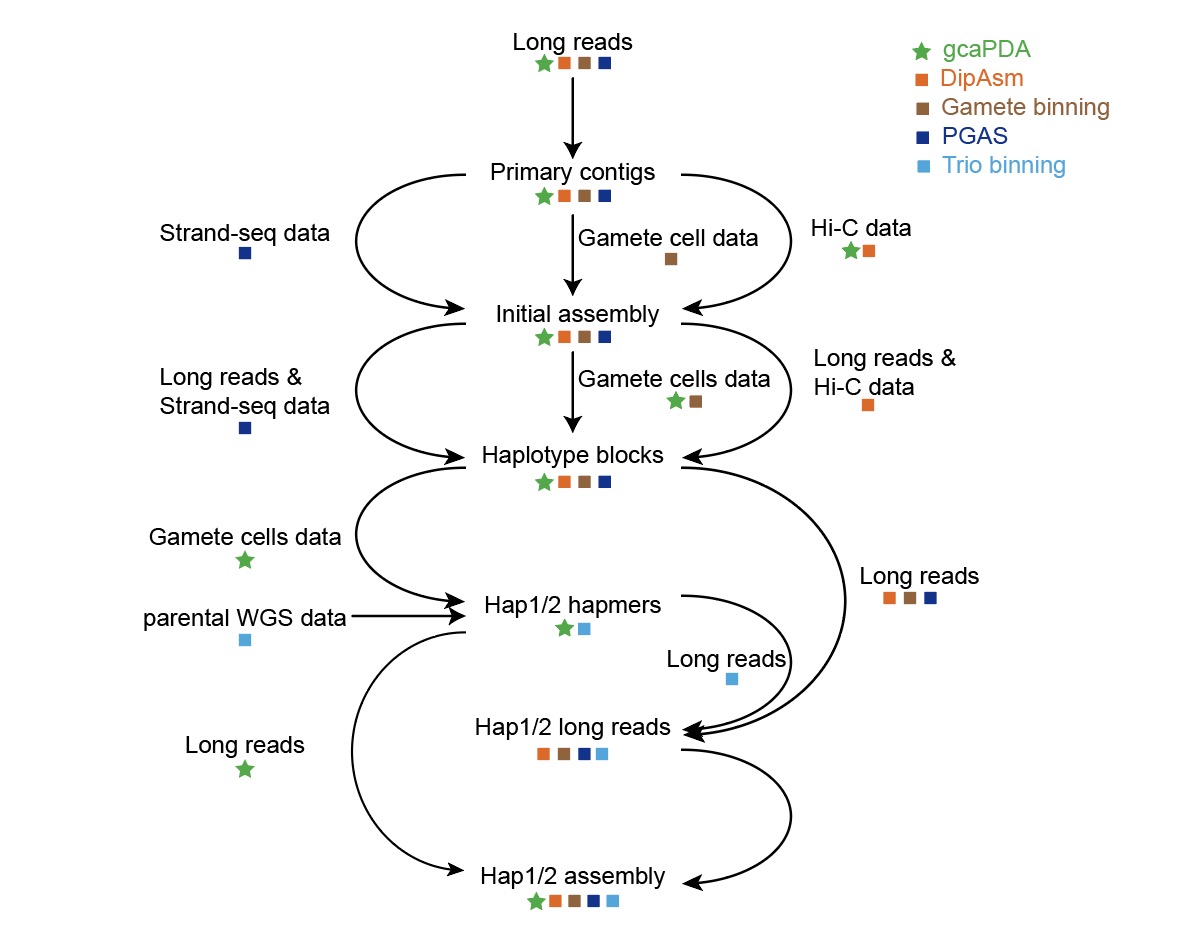

**Supplementary Figure 1.** **Comparison of framework of different diploid assemblers.** DipAsm, PGAS, Gamete-binning and gcaPDA starts with assembling long reads into primary contigs and scaffolding contigs into superscaffolds (initial assembly). Reads data was then mapped to the initial assembly to identify and phase variations into haplotype blocks. In DipAsm, gamete binning and PGAS, long reads were partitioned into haplotypes based on variations and each haplotype was assembled from partitioned long reads, respectively. In trio binning method, long reads were partitioned into haplotypes based on hapmers derived from parental WGS data and each haplotype was assembled from partitioned long reads, respectively. In contrast, gcaPDA partitioned gamete cell reads based on haplotype blocks to and generated hapmers. In gcaPDA, unpartitioned long reads and hapmers were used to assemble both haplotypes simultaneously. Data and analysis steps used in gcaPDA were indicated by green stars, while data and analysis steps used in DipAsm, gamete binning, PGAS and trio binning were indicated by orange, brown, deep blue and light blue cubes, respectively.

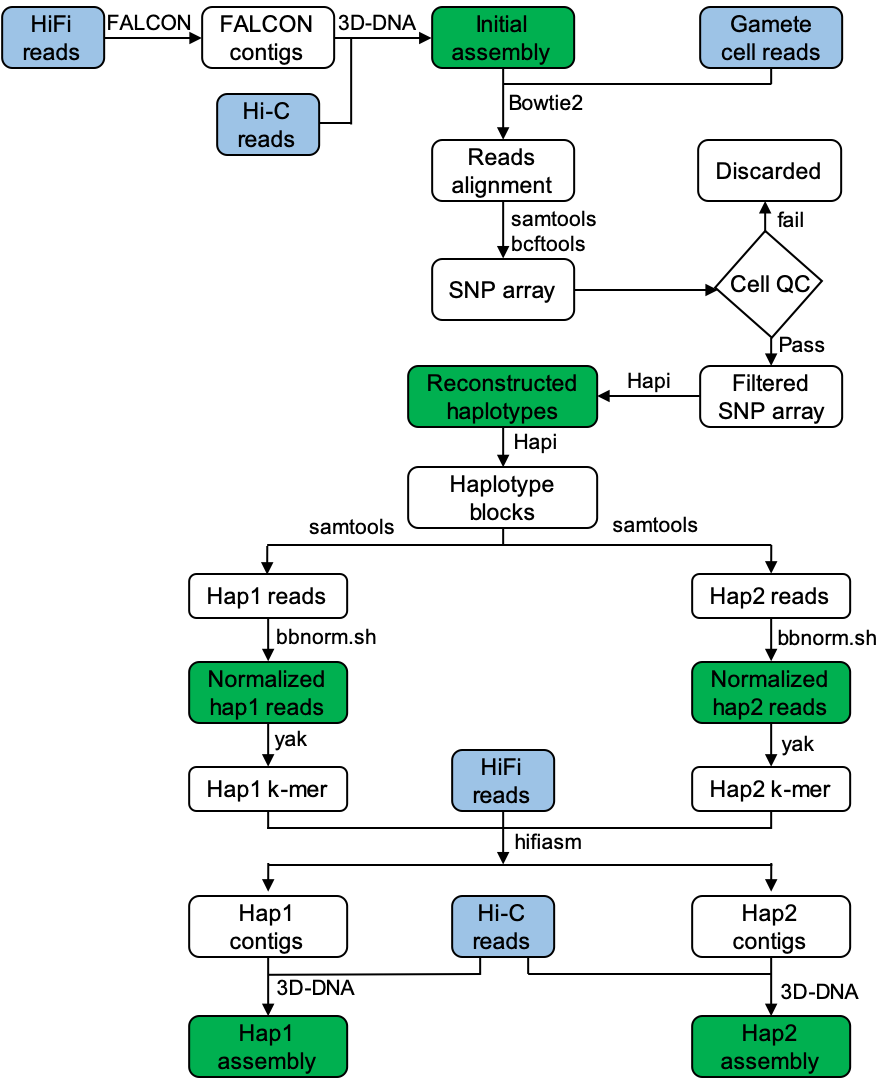

**Supplementary Figure 2**. **The analysis workflow of gcaPDA.** gcaPDA consists of 4 major steps: 1) building an initial assembly; 2) reconstruction of haplotypes; 3) partition and normalization of gamete cell reads and 4) generating chromosome-scale phased diploid assembly. Data are shown in round rectangles. Input data set (HiFi reads, Hi-C reads, gamete cell reads) highlighted in blue and result of each major step is highlighted in green. Software used at each analysis step is shown on the top/right.

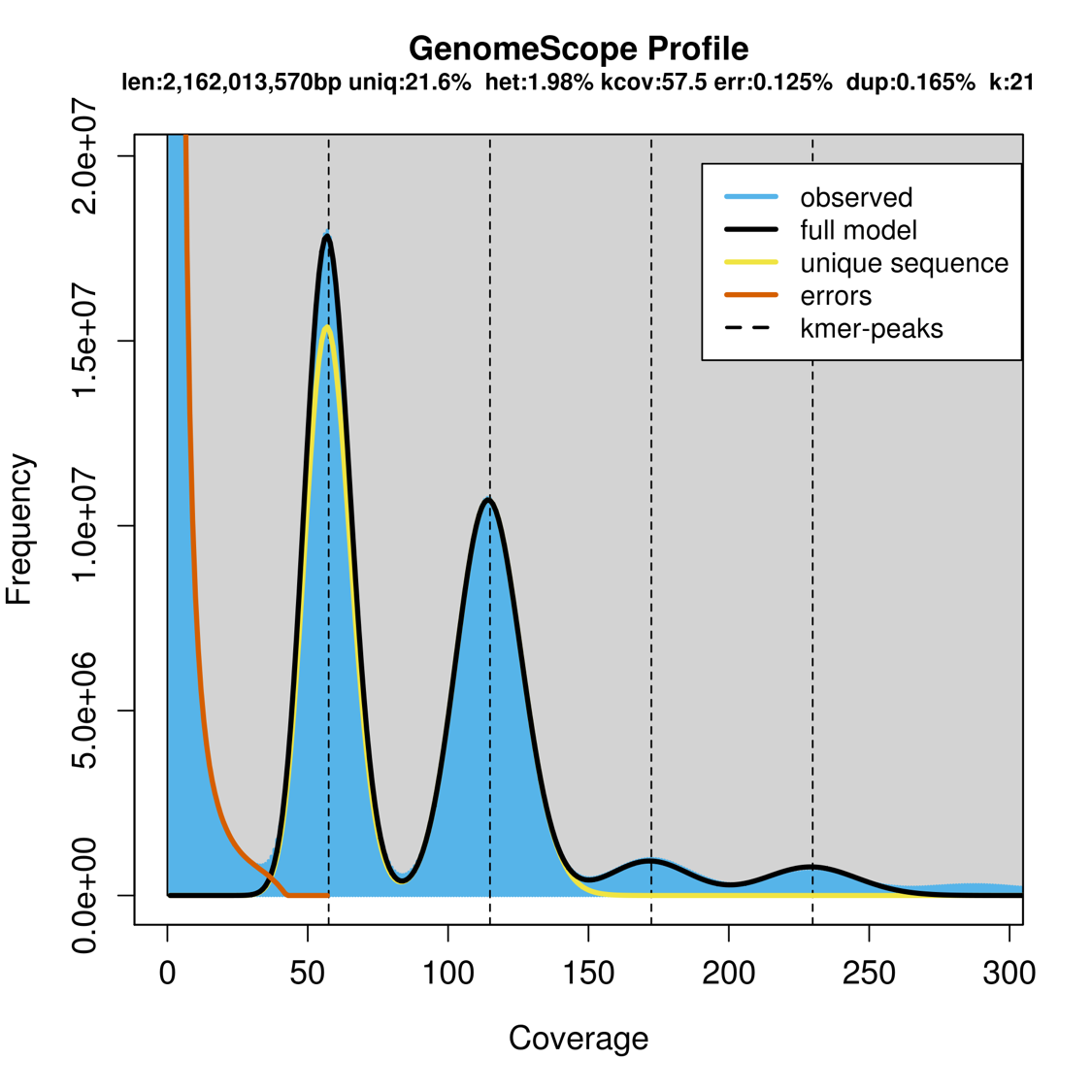

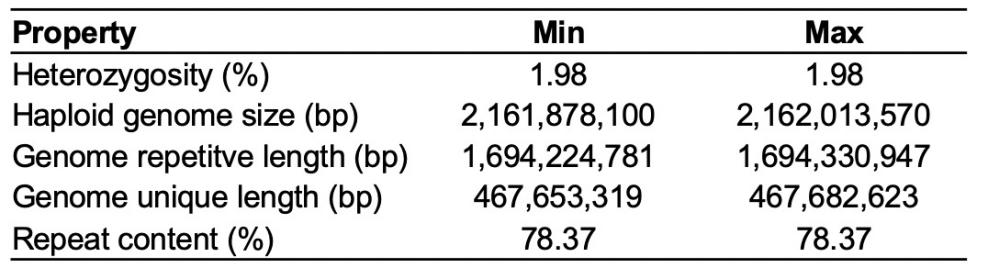

**Supplementary Figure 3.** **Genome survey of the SK x B73 F_1_ hybrid.** Simulated HiFi reads were used for this analysis (k-mer size =21). Haploid genome size and heterozygosity of F_1_ hybrid was estimated as 2.16 Gb and 1.98%, respectively.

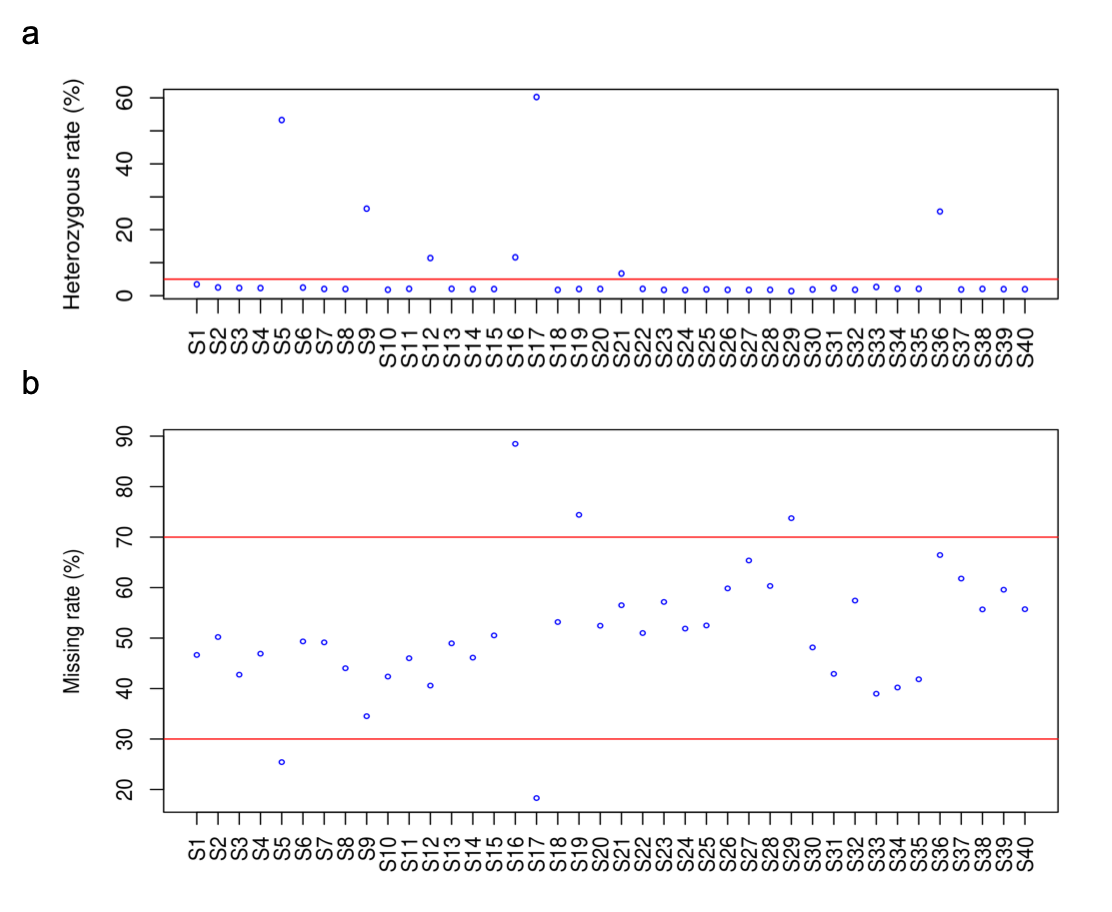

**Supplementary Figure 4**. **Quality control of gamete cells based on SNP statistics**. **a**), heterozygous rate of SNPs of gamete cells. Gamete cell with heterozygous rate higher that 5% (cutoff indicated by red line) were considered as contaminated and excluded from downstream analysis. **b**), SNP missing rate of gamete cells. Gamete cells with SNP missing rate >70% or <30% were considered as low-quality cells and excluded from downstream analysis.

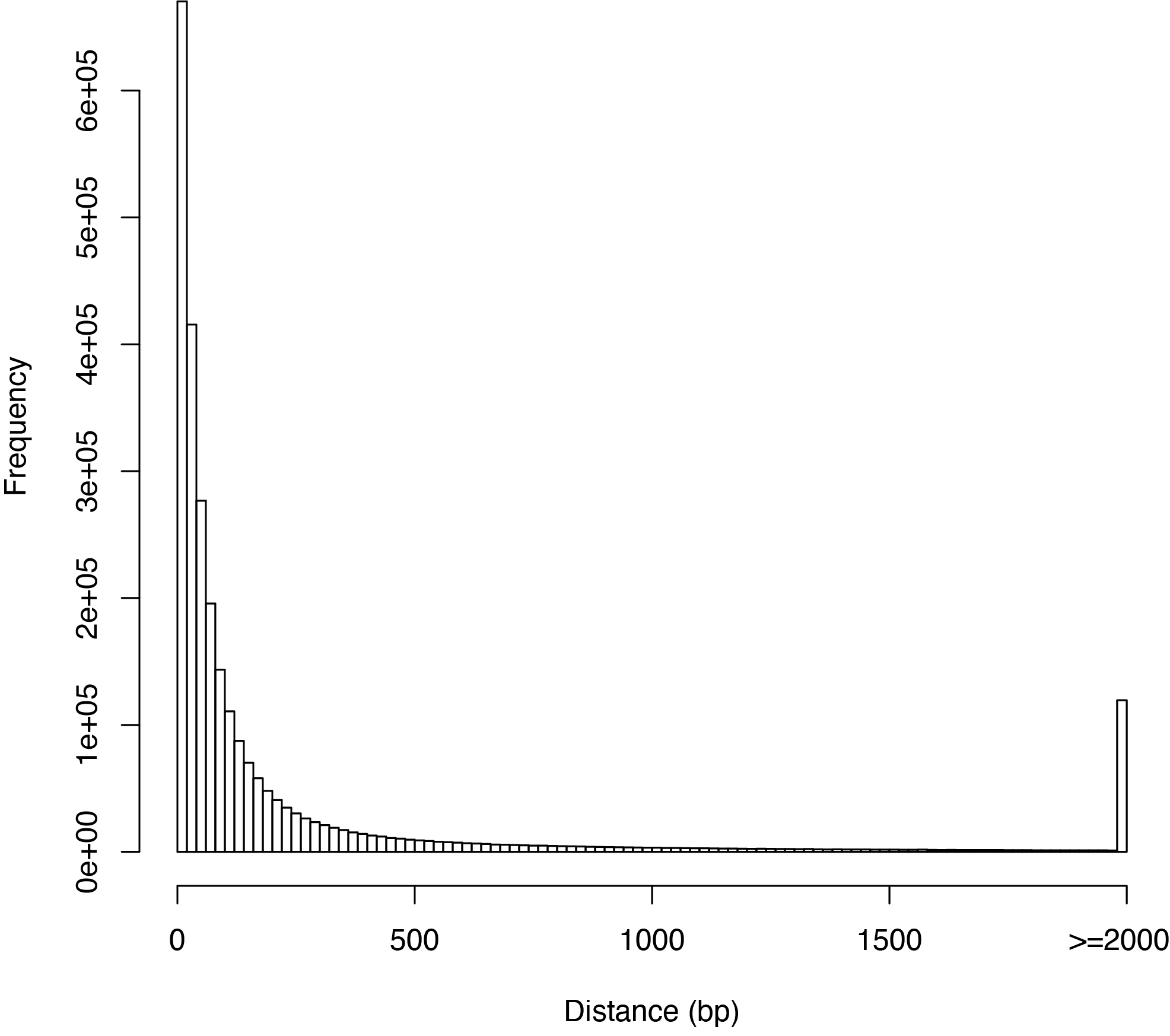

**Supplementary Figure 5.** **Distribution of distance between adjacent SNPs in the reconstructed haplotypes.**

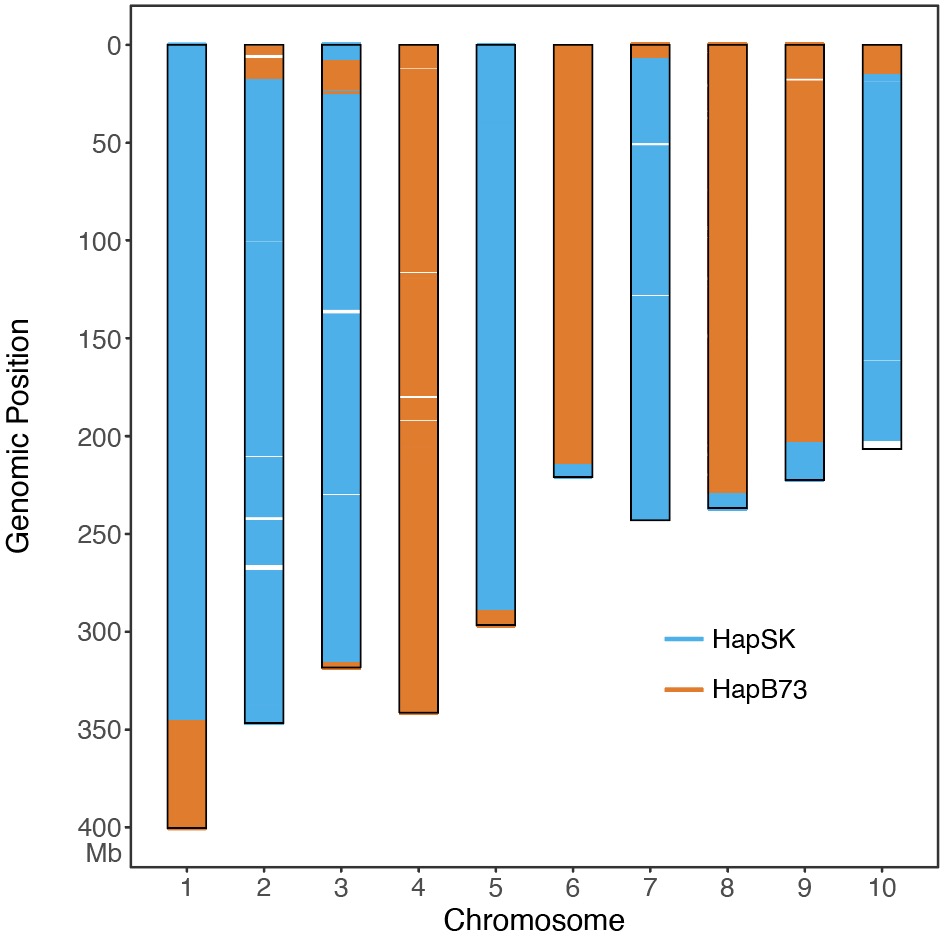

**Supplementary Figure 6.** **Visualization of haplotype blocks of gamete cell S2.** Chromosomes of the initial assembly was depicted with black rectangles. The genotype of gamete cell S2 at each SNP locus was compared with the reconstructed haplotype. SNP loci with identical genotype to hapSK were depicted with blue line, while SNP loci with identical genotype to hapB73 were depicted with orange line.

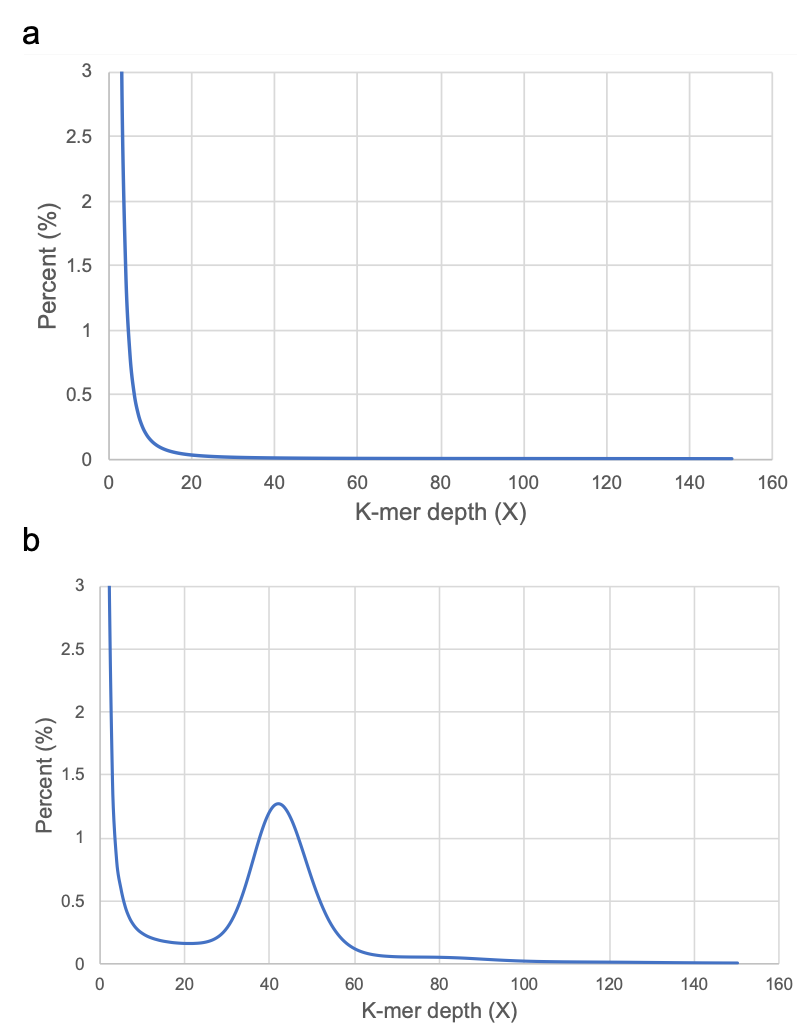

**Supplementary Figure 7**. K-mer distribution of haplotype reads before **a**) and after normalization **b**). We used k-mer size of length 31 for this analysis.

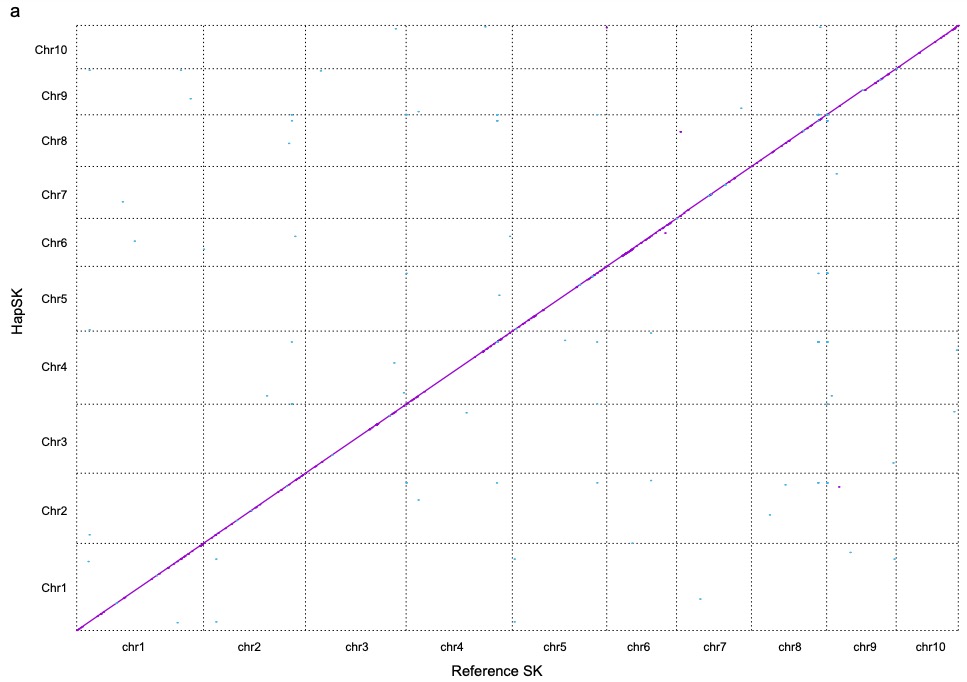

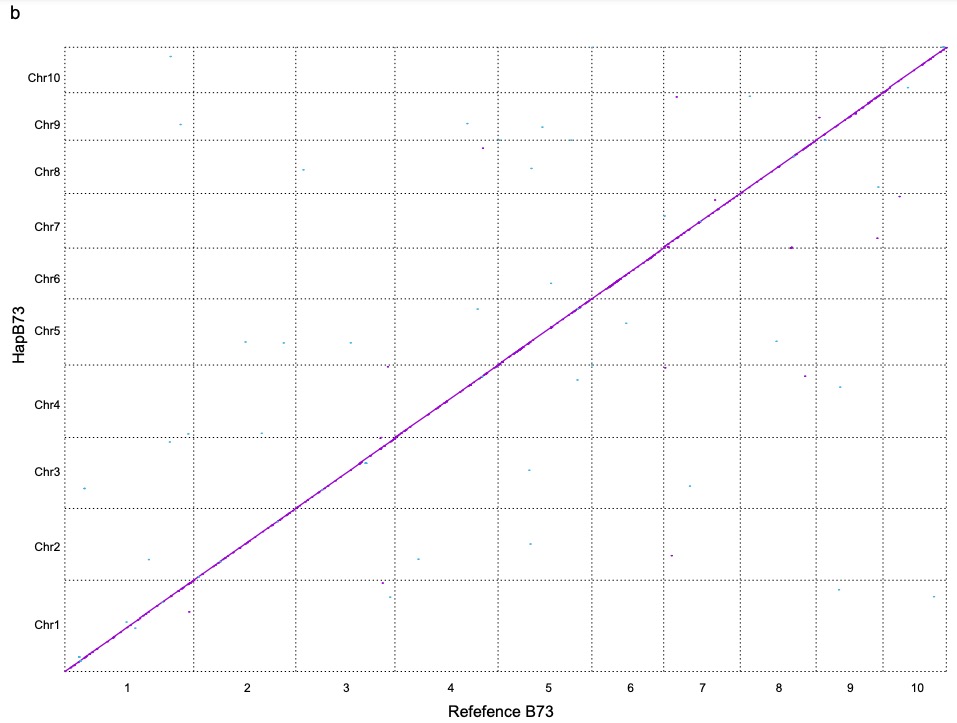

**Supplementary Figure 8.** **Visualization of sequence alignments between gcaDPA assembly and parental genomes.** a) Visualization of alignments between hapSK chromosomes and reference SK chromosomes. b) Visualization of alignments between hapB73 chromosomes and reference B73 chromosomes.

K-mer (%)

| No. of cells | SK | B73 | SK +B73 | Simulation |
| --- | --- | --- | --- | --- |
| 1 | 55.3503 | 62.2272 | 52.4046 | 50 |
| 2 | 77.906 | 79.6487 | 73.1091 | 75 |
| 3 | 85.3579 | 88.5841 | 82.4131 | 87.5 |
| 4 | 90.7209 | 92.4858 | 88.1911 | 93.75 |
| 5 | 95.1403 | 95.2358 | 93.0872 | 96.875 |
| 6 | 96.6041 | 96.8826 | 95.2552 | 98.4375 |
| 7 | 97.2606 | 97.6033 | 96.2397 | 99.2188 |
| 8 | 97.8698 | 98.0324 | 96.992 | 99.6094 |
| 9 | 98.2235 | 98.236 | 97.4021 | 99.8047 |
| 10 | 98.5648 | 98.4371 | 97.8 | 99.9023 |
| 11 | 98.7171 | 98.5256 | 97.9765 | 99.9512 |
| 12 | 98.8986 | 98.6046 | 98.1655 | 99.9756 |
| 13 | 98.954 | 98.6887 | 98.2682 | 99.9878 |
| 14 | 99.0442 | 98.7755 | 98.3987 | 99.9939 |
| 15 | 99.0974 | 98.8408 | 98.4848 | 99.9969 |
| 16 | 99.1476 | 98.8784 | 98.5485 | 99.9985 |
| 17 | 99.1776 | 98.9054 | 98.5897 | 99.9992 |
| 18 | 99.192 | 98.921 | 98.6109 | 99.9996 |
| 19 | 99.2056 | 98.9394 | 98.634 | 99.9998 |
| 20 | 99.2165 | 98.9521 | 98.651 | 99.9999 |
| 21 | 99.2289 | 98.9704 | 98.6725 | 100 |
| 22 | 99.2387 | 98.9855 | 98.6896 | 100 |
| 23 | 99.2462 | 98.991 | 98.6987 | 100 |
| 24 | 99.2531 | 98.999 | 98.7089 | 100 |
| 25 | 99.2598 | 99.0058 | 98.7182 | 100 |
| 26 | 99.2655 | 99.0125 | 98.7271 | 100 |
| 27 | 99.2701 | 99.0183 | 98.7346 | 100 |
| 28 | 99.2754 | 99.0246 | 98.7427 | 100 |
| 29 | 99.2816 | 99.0297 | 98.7506 | 100 |
| 30 | 99.2857 | 99.0335 | 98.756 | 100 |
| 31 | 99.2897 | 99.0387 | 98.7626 | 100 |

**Supplementary Figure 9.** **Accumulated percentage of genomic k-mers covered in gamete cells reads**. Genomic k-mers set were calculated from SK, B73 genome, or both (SK+B73). A genomic k-mer was defined was covered if its frequency in gamete cell reads is >4. A simulated accumulation curve was also plotted, assuming that each gamete cell randomly covered 50% of genomic k-mers.

**Supplementary Tables**

**Supplementary Table 1.** Statistics of simulated reads.

| **Simulated data set** | **Error rate** | **Read No.** | **Total read length (bp)** | **Average read length (bp)** |
| --- | --- | --- | --- | --- |
| B73 HiFi reads | 0.002 | 9,042,308 | 126,261,011,397 | 13,963 |
| SK HiFi reads | 0.002 | 9,255,246 | 129,233,929,145 | 13,963 |
| B73 short reads | 0.01 | 600,000,000 | 90,000,000,000 | 150 |
| SK short reads | 0.01 | 600,000,000 | 90,000,000,000 | 150 |

**Supplementary Table 2**. Statistics of Hi-C sequencing reads.

| **Tissue** | **Total read pair** | **Total read length (bp)** |
| --- | --- | --- |
| Roots | 1,423,190,273 | 426,957,081,900 |

**Supplementary Table 3.** Statistics of sequencing reads of gamete cells.

| Sample | Raw reads | | Clean reads | |
| --- | --- | --- | --- | --- |
|  | Count (million) | Read base (Gb) | Count (million) | Read base (Gb) |
| S1 | 399.1 | 59.9 | 375.1 | 55.2 |
| S2 | 415.2 | 62.3 | 389.8 | 57.4 |
| S3 | 417.6 | 62.6 | 401.8 | 59.4 |
| S4 | 400.1 | 60.0 | 384.6 | 56.8 |
| S5 | 393.3 | 59.0 | 379.4 | 55.9 |
| S6 | 415.1 | 62.3 | 401.5 | 59.2 |
| S7 | 500.7 | 75.1 | 488.9 | 72.5 |
| S8 | 494.2 | 74.1 | 482.8 | 71.6 |
| S9 | 523.0 | 78.5 | 510.8 | 75.7 |
| S10 | 523.9 | 78.6 | 512.4 | 76.0 |
| S11 | 493.3 | 74.0 | 471.6 | 69.6 |
| S12 | 518.8 | 77.8 | 497.9 | 73.5 |
| S13 | 369.4 | 55.4 | 344.2 | 50.5 |
| S14 | 380.7 | 57.1 | 355.4 | 52.2 |
| S15 | 345.3 | 51.8 | 321.3 | 47.2 |
| S16 | 338.0 | 50.7 | 317.1 | 46.6 |
| S17 | 508.9 | 76.3 | 492.2 | 72.7 |
| S18 | 501.0 | 75.2 | 487.3 | 72.1 |
| S19 | 457.6 | 68.6 | 443.2 | 65.5 |
| S20 | 483.9 | 72.6 | 469.1 | 69.3 |
| S21 | 437.0 | 65.5 | 421.3 | 62.1 |
| S22 | 382.9 | 57.4 | 368.3 | 54.3 |
| S23 | 376.6 | 56.5 | 364.2 | 53.7 |
| S24 | 443.0 | 66.5 | 429.7 | 63.5 |
| S25 | 394.1 | 59.1 | 379.6 | 56.1 |
| S26 | 326.3 | 48.9 | 314.0 | 46.4 |
| S27 | 482.0 | 72.3 | 468.0 | 69.3 |
| S28 | 371.4 | 55.7 | 359.8 | 53.3 |
| S29 | 429.3 | 64.4 | 415.6 | 61.5 |
| S30 | 413.6 | 62.0 | 398.6 | 58.8 |
| S31 | 413.8 | 62.1 | 400.5 | 59.2 |
| S32 | 463.6 | 69.5 | 449.3 | 66.5 |
| S33 | 289.0 | 43.4 | 270.9 | 39.6 |
| S34 | 366.5 | 55.0 | 346.7 | 50.7 |
| S35 | 307.9 | 46.2 | 289.1 | 42.3 |
| S36 | 337.8 | 50.7 | 319.1 | 46.7 |
| S37 | 404.7 | 60.7 | 392.1 | 58.0 |
| S38 | 463.8 | 69.6 | 448.8 | 66.3 |
| S39 | 393.2 | 59.0 | 379.1 | 56.0 |
| S40 | 455.8 | 68.4 | 438.4 | 64.7 |

**Supplementary Table 4**. Purge haplotigs from FALCON primary contigs.

| **Contigs** | **Primary contigs** | **Artefacts** | **Haplotigs** | **Clean contigs** |
| --- | --- | --- | --- | --- |
| Contig No. | 5,014 | 141 | 2,462 | 2,411 |
| Contig N50 (bp) | 1,707,428 | 24,630 | 357,307 | 2,017,689 |
| Contig length (bp) | 2,903,282,308 | 2,896,924 | 428,970,525 | 2,471,414,859 |

**Supplementary Table 5**. Statistics of SNPs identified in gamete cells.

| Sample | Homozygous calls | Heterozygous calls | Missing calls | Heterozygous rate (%) | Missing rate (%) |
| --- | --- | --- | --- | --- | --- |
| S1 | 2,100,669 | 74,653 | 1,902,968 | 3.43 | 46.66 |
| S2 | 1,980,218 | 50,490 | 2,047,582 | 2.49 | 50.21 |
| S3 | 2,280,119 | 54,649 | 1,743,522 | 2.34 | 42.75 |
| S4 | 2,115,073 | 49,978 | 1,913,239 | 2.31 | 46.91 |
| S5^*^ | 1,421,017 | 1,620,736 | 1,036,537 | 53.28 | 25.42 |
| S6 | 2,015,379 | 50,726 | 2,012,185 | 2.46 | 49.34 |
| S7 | 2,031,329 | 42,354 | 2,004,607 | 2.04 | 49.15 |
| S8 | 2,236,783 | 46,034 | 1,795,473 | 2.02 | 44.03 |
| S9^*^ | 1,964,351 | 705,800 | 1,408,139 | 26.43 | 34.53 |
| S10 | 2,307,611 | 42,321 | 1,728,358 | 1.80 | 42.38 |
| S11 | 2,155,980 | 45,819 | 1,876,491 | 2.08 | 46.01 |
| S12^*^ | 2,146,651 | 276,147 | 1,655,492 | 11.40 | 40.59 |
| S13 | 2,038,436 | 43,255 | 1,996,599 | 2.08 | 48.96 |
| S14 | 2,153,800 | 43,283 | 1,881,207 | 1.97 | 46.13 |
| S15 | 1,977,077 | 40,646 | 2,060,567 | 2.01 | 50.53 |
| S16^*^ | 415,674 | 54,632 | 3,607,984 | 11.62 | 88.47 |
| S17^*^ | 1,324,847 | 2,007,100 | 746,343 | 60.24 | 18.30 |
| S18 | 1,875,931 | 33,704 | 2,168,655 | 1.76 | 53.18 |
| S19^*^ | 1,022,559 | 20,965 | 3,034,766 | 2.01 | 74.41 |
| S20 | 1,899,502 | 39,603 | 2,139,185 | 2.04 | 52.45 |
| S21^*^ | 1,654,560 | 119,133 | 2,304,597 | 6.72 | 56.51 |
| S22 | 1,957,133 | 41,323 | 2,079,834 | 2.07 | 51.00 |
| S23 | 1,716,481 | 30,664 | 2,331,145 | 1.76 | 57.16 |
| S24 | 1,928,832 | 33,566 | 2,115,892 | 1.71 | 51.88 |
| S25 | 1,900,530 | 36,837 | 2,140,923 | 1.90 | 52.50 |
| S26 | 1,608,829 | 29,059 | 2,440,402 | 1.77 | 59.84 |
| S27 | 1,387,594 | 24,601 | 2,666,095 | 1.74 | 65.37 |
| S28 | 1,589,609 | 28,721 | 2,459,960 | 1.77 | 60.32 |
| S29^*^ | 1,055,719 | 14,869 | 3,007,702 | 1.39 | 73.75 |
| S30 | 2,074,821 | 39,686 | 1,963,783 | 1.88 | 48.15 |
| S31 | 2,275,225 | 53,102 | 1,749,963 | 2.28 | 42.91 |
| S32 | 1,704,902 | 31,292 | 2,342,096 | 1.80 | 57.43 |
| S33 | 2,423,031 | 65,624 | 1,589,635 | 2.64 | 38.98 |
| S34 | 2,386,849 | 51,447 | 1,639,994 | 2.11 | 40.21 |
| S35 | 2,322,732 | 49,391 | 1,706,167 | 2.08 | 41.84 |
| S36^*^ | 1,018,799 | 349,340 | 2,710,151 | 25.53 | 66.45 |
| S37 | 1,529,145 | 29,355 | 2,519,790 | 1.88 | 61.79 |
| S38 | 1,770,185 | 37,122 | 2,270,983 | 2.05 | 55.68 |
| S39 | 1,615,903 | 32,695 | 2,429,692 | 1.98 | 59.58 |
| S40 | 1,770,442 | 35,391 | 2,272,457 | 1.96 | 55.72 |

^*^The nine gamete cells that failed quality control were marked with asterisk.

**Supplementary Table 6.** Statistics of k-mer in the parental reference genomes.

| **Category** | **Distinct k-mer count** | **Percent (%)** |
| --- | --- | --- |
| Union (B73, SK) | 799,441,609 | 100.0 |
| B73 | 607,325,429 | 76.0 |
| SK | 608,801,647 | 76.2 |
| Intersection (B73, SK) | 416,685,467 | 52.1 |
| B73 hapmer | 190,639,962 | 23.8 |
| SK hapmer | 192,116,180 | 24.0 |

**Supplementary Table 7.** Statistics of whole genome sequence comparison between haplotype assemblies and reference assemblies.

| **Comparison** | **HapSK vs SK reference** | | **HapB73 vs B73 reference** | |
| --- | --- | --- | --- | --- |
|  | **HapSK** | **SK reference** | **HapB73** | **B73 reference** |
| Total bases (bp) | 2,162,128,026 | 2,153,898,797 | 2,158,996,842 | 2,104,350,182 |
| Alignment coverage (%) | 99.03 | 99.04 | 98.05 | 99.72 |
| 1-1 alignment identity (%) | 99.99 | 99.99 | 99.99 | 99.99 |
| Relocations | 28 | 28 | 20 | 45 |
| Translocations | 81 | 47 | 167 | 50 |
| Inversions | 6 | 5 | 3 | 5 |
| Insertions (>1bp) | 1,822 | 2,051 | 2,848 | 963 |
| SNPs | 17,132 | 17,132 | 19,830 | 19,830 |
| Indels (1bp) | 22,560 | 22,560 | 36,695 | 36,695 |

Supplementary Table 8. Statistics of haplotype blocks from the other haplotype.

|  | **Chromosomes** | | **Un-anchored sequences** | |
| --- | --- | --- | --- | --- |
|  | **Length (bp)** | **Percent (%)** | **Length (bp)** | **Percent (%)** |
| HapSK assembly | 2,086,840,877 | 96.52 | 75,287,149 | 3.48 |
| HapB73 assenbly | 2,104,568,114 | 97.48 | 54,428,728 | 2.52 |
| HapB73 blocks in HapSK assembly | 22,445,582 | 1.08 | 22,468,175 | 29.84 |
| HapSK blocks in HapB73 assembly | 31,273,932 | 1.49 | 42,870,239 | 78.76 |
| Total mis-assigned blocks | 53,719,514 | 1.28 | 65,338,414 | 50.37 |
